# Variant-specific Mendelian Risk Prediction Model

**DOI:** 10.1101/2023.03.06.531363

**Authors:** Eunchan Bae, Julie-Alexia Dias, Theodore Huang, Jinbo Chen, Giovanni Parmigiani, Timothy R. Rebbeck, Danielle Braun

**Author notes:** **Correspondence** Danielle Braun, Department of Data Sciences, Dana-Farber Cancer Institute, 450 Brookline Ave, CLSB 11051, Boston, MA 02215, USA. Eunchan Bae, Department of Biostatistics, Epidemiology and Informatics, University of Pennsylvania, Philadelphia, PA, USA. **Funding information** NCI, Grant/Award Number: R01-CA207365-01A1 and 5P30CA006516-54.

## Abstract

Many pathogenic sequence variants (PSVs) have been associated with increased risk of cancers. Mendelian risk prediction models use Mendelian laws of inheritance to predict the probability of having a PSV based on family history, as well as specified PSV frequency and penetrance (agespecific probability of developing cancer given genotype). Most existing models assume penetrance is the same for any PSVs in a certain gene. However, for some genes (for example, BRCA1/2), cancer risk does vary by PSV. We propose an extension of Mendelian risk prediction models to relax the assumption that risk is the same for any PSVs in a certain gene by incorporating variant-specific penetrances and illustrating these extensions on two existing Mendelian risk prediction models, BRCAPRO and PanelPRO. Our proposed BRCAPRO-variant and PanelPRO-variant models incorporate variant-specific BRCA1/2 PSVs through the region classifications. Due to the sparsity of the variant information we classify BRCA1/2 PSVs into three regions; the breast cancer clustering region (BCCR), the ovarian cancer clustering region (OCCR), and an other region. Simulations were conducted to evaluate the performance of the proposed BRCAPRO-variant model compared to the existing BRCAPRO model which assumes the penetrance is the same for any PSVs in BRCA1 (and respectively BRCA2). Simulation results showed that the BRCAPRO-variant model was well calibrated to predict region-specific BRCA1/2 carrier status with high discrimination and accuracy on the region-specific level. In addition, we showed that the BRCAPRO-variant model achieved performance gains over the existing risk prediction models in terms of calibration without loss in discrimination and accuracy. We also evaluated the performance of the two proposed models, BRCAPRO-variant and PanelPRO-variant, on a cohort of 1,961 families from the Cancer Genetics Network (CGN). We showed that our proposed models provide region-specific PSV carrier probabilities with high accuracy, while the calibration, discrimination and accuracy of gene-specific PSV carrier probabilities were comparable to the existing gene-specific models. As more variant-specific PSV penetrances become available, we have shown that Mendelian risk prediction models can be extended to integrate the additional information, providing precise variant or region-specific PSV carrier probabilities and improving future cancer risk predictions.

## 1 INTRODUCTION

A number of genes have been identified for which inherited pathogenic sequence variants (PSVs) are significantly associated with cancer risk [1]. Some PSVs are associated with cancer syndromes in which phenotypes cluster in high risk families. According to National Cancer Institute, cancer caused by PSVs accounts for about 5% to 10% of all cancers [2].

With development of DNA sequencing techniques [3], high-risk individuals are increasingly referred for multigene panel genetic testing to assess inherited genetic susceptibility [4, 5]. Clinicians are using germline testing results to identify individuals at high risk of developing cancer and apply appropriate preventive interventions, including risk-reducing surgical or chemopreventive procedures [6]. Carrier probability assessment facilitates the task of identifying individuals who are likely to harbor PSVs and is a key step in precision prevention.

Two approaches have been commonly used in carrier probability assessment. *Empirical* approaches model the conditional distribution of genotype given family history phenotypes as covariates, for example using logistic regression. In contrast, *Mendelian* models are constructed using the conditional distributions of phenotypes given genotypes (penetrance), and the prior distributions of ethnicity-specific genotypes (prevalence). They then use Bayes theorem and Mendelian law of inheritance to calculate the posterior probability of genotype carrier status and future risk of cancers [7, 8, 9]. These models are widely used clinically [10, 11, 12]. Some of these models are able to incorporate additional information such as the occurrence of second cancers [13, 14].

There are several existing Mendelian models to predict any variant-specific PSV carrier status which assumes the risk is the same across all PSVs at a locus, such as PanelPRO for 22 genes including BRCA1 and BRCA2, BOADICEA [14] for the BRCA1, BRCA2, PALB2, CHEK2, and ATM genes, MMRPRO for MMR genes [15], PancPRO for a hypothetical “PANC” gene [16], MelaPRO for the CDKN2A gene [17] and BRCAPRO [18] for the BRCA1 and BRCA2 genes. In particular, BRCAPRO [19] assumes BRCA1 (and respectively BRCA2) penetrances are the same across all PSV in BRCA1 (and respectively BRCA2). However, it is known that risk for several cancers differs by PSV within a gene. For example, in E-cadherin (CDH1), Lo et al. [9] showed that individuals with PSVs in the PRE-PRO region are six times more likely to have family members with colorectal cancer compared to those with PSVs in other regions. Southey et al [20] showed that among European women, the magnitude of increased risk of breast cancer varies by PSV among PALB2 and CHEK2 carriers: c.1592delT (OR: 4.52) and c.3113G>A (OR: 5.93) in PALB2 carriers and c.349A>G (OR: 2.26), c.1036C>T (OR: 5.06), and c.538C>T (OR: 1.33) in CHEK2 carriers. For PSVs in BRCA1 and BRCA2, Rebbeck et al. [21] reported that breast and ovarian cancer risks differ by PSV or domains in which groups of PSVs are found among female carriers. Patel et al. [22] showed that PSVs in c.7914+ in BRCA2 male carriers were significantly associated with elevated risk of prostate cancer compared with PSVs in the reference bin c.1001-c.7913 (HR:1.78) and elevated risk of Gleason 8+ prostate cancer (HR: 3.11). Patel et al. also showed that PSVs in c.756-c.1000 in BRCA2 male carriers were also associated with elevated prostate cancer risk (HR: 2.83) and elevated risk of Gleason 8+ prostate cancer (HR: 4.95).

As cancer risk can vary across different PSVs in the same gene, it is important to extend the existing models to estimate variant-specific PSV carrier probabilities and future cancer risk. In this work, we introduce a general framework for variant-specific Mendelian risk prediction models. Section 2 reviews existing Mendelian models and extends them to incorporate variant-specific PSV penetrances. Section 3 presents the simulation studies and Section 4 presents an application to the Cancer Genetics Network cohort data in the context of BRCA1/2.

## 2 METHOD

### 2.1 Mendelian risk prediction models

Mendelian risk prediction models predict the probability of carrying PSVs in a range of genes based on family history and are trained using prior knowledge of prevalences of PSVs in these genes. Family history consists of the age at which each family member was diagnosed with cancer, or the unaffected family member’s current age (censoring age). Suppose there are *R* types of cancers that are associated with a PSV found in a family of *n* members. Let *i* index the family member, and let *i* =1 indicate the proband. Let *T*_*ri*_ be the diagnosis age of the *r* -th cancer for the *i* -th family member, and let *C*_*i*_ indicate the censoring age. Let *X*_*ri*_ = min(*T*_*ri*_, *C*_*i*_) indicate the observed event age for the *r* -th cancer, and *δ*_*ri*_ = I(*T*_*ri*_ ≤ *C*_*i*_). We denote the individual history for the *i* -th family member by **H**_*i*_ = (*X*_1*i*_, …, *X*_*Ri*_, *δ*_1*i*_, …, *δ*_*Ri*_) and the overall family history **H** = (**H**_1_, …, **H**_*n*_). In addition to the family history, let *U*_*i*_ be the indicator of the *i* -th individual being male and **U** = (*U*_1_, …, *U*_*n*_). Suppose in addition there are *J* individual-level covariates and let **Y**_*i*_ = (*Y*_1*i*_, *Y*_2*i*_, …, *Y*_*Ji*_) be the *i* -th family member’s covariates (e.g., ethnicity, or history of medical preventative interventions). Let **Y** = (**Y**_1_, …, **Y**_*n*_) be the collection of covariates. Suppose there are *M* genes of interest and let *G*_*mi*_ denote the PSV carrier status for individual *i* for gene *m* (*G*_*mi*_ = 1 if individual *i* has a PSV in gene *m* and 0 otherwise). *G*_*mi*_ = 1 for an autosomal dominant gene *m* when family member *i* carries at least one deleterious mutant allele, or both deleterious alleles for an autosomal recessive gene. Generalization to variant-specific analyses proceeds by allowing *G*_*mi*_ to be a multinomial variable. The extended variant-specific Mendelian risk prediction model is introduced in Section 2.2. Here we assume all PSVs within a gene have the same risk, and assume a binary variable for *G*_*mi*_, this is relaxed in the following section. Let **G**_*i*_ = (*G*_1*i*_, …, *G*_*M i*_) be the genotype of interest for the *i* -th family member and **G** = (**G**_1_, …, **G**_*n*_) be the collection of genotypes for the family.

Using Bayes theorem we can calculate P(**G**_1_|**H, Y, U**), the probability of the counselee having a PSV given family history, sex and individual-specific covariates. For simplicity we will omit **U** and **Y** from subsequent equations.

Using Bayes’ theorem,

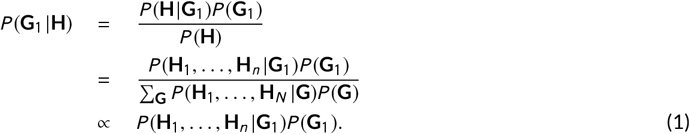

The prior *P* (**G**_1_) (=*P* (**G**_1_ |**U, Y**) =*P* (**G**_1_ |**Y**)) is the population-level probability of genotypes and is ethnicity-specific and not sex-specific, and *P* (**H**_1_, …, **H**_*n*_ |**G**_1_) can be calculated as follows:

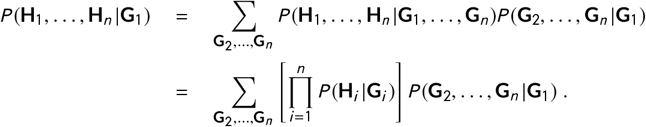

The last equality assumes each family member’s cancer history is conditionally independent given the genotype. Relaxation of this assumption is discussed by Huang et al. [23]. Assuming Hardy-Weinberg equilibrium [24], *P* (**G**_2_, …, **G**_*n*_ |**G**_1_) can be calculated based on Mendelian laws of inheritance, and subsequently marginalized depending on which gene is considered dominant or recessive. *P* (**H**_*i*_ |**G**_*i*_) is derived from the penetrance functions.

Therefore, Mendelian risk prediction models calculate the posterior probability of the counselee having a PSV as

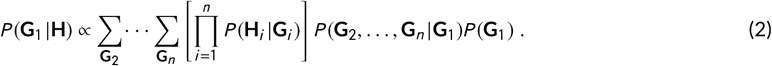

### 2.2 Variant-specific Mendelian Risk Prediction Models

We now describe how to extend this approach to allow for variant-specific risk. Let **G**_1_ indicate the counselee’s variant-specific PSV status of interest and **G**_1_ = (*G*_11_, *G*_21_, …, *G*_*M* 1_). As defined earlier, let *G*_*mi*_ indicate the PSV in gene *m* for individual *i*. Let *G*_*mi*_ = *g*_*mi*_ indicate that individual *i* carries PSV *g*_*mi*_ in gene *m*, and *G*_*mi*_ = 0 indicate that individual *i* carriers no PSV in gene *m* (note, this is no longer the binary indicator specified earlier, as there are multiple PSVs for a gene). The proposed variant-specific Mendelian risk prediction model calculates the variant-specific probabilities for *m* genes by expanding Equation (1) as follows,

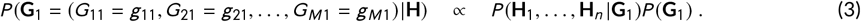

In Equation (3), the prior *P* (**G**_1_)(=*P* (**G**_1_ |**U, Y**) =*P* (**G**_1_ |**Y**)) is the probability of variant-specific genotypes and is ethnicity-specific, and not sex-specific. To calculate *P* (**H** |**G**_1_) = *P* (**H**_1_, …, **H**_*n*_ |**G**_1_) we assume, as earlier, that histories of each family member are conditionally independent given the variant-specific genotype. *P* (**H**_*i*_ |**G**_*i*_) is derived from the variant-specific PSV penetrance functions and estimated from published studies, for example, see [25].

Assuming Hardy-Weinberg equilibrium, *P* (**G**_2_, …, **G**_*n*_ |**G**_1_) can be calculated based on Mendelian law of inheritance. The posterior probability of the counselee having PSVs be written as;

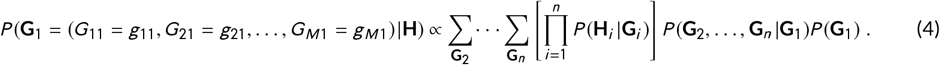

### 2.3 Model Parameters

In variant-specific Mendelian risk prediction models, we need to estimate the prevalences of PSVs, *P* (**G**_1_ = (*G*_11_ = *g*_11_, *G*_21_ = *g*_21_, …, *G*_*M* 1_ = *g*_*M* 1_), as well as the variant-specific penetrances, *P* (**H** |**G**_1_ = (*G*_11_ = *g*_11_, *G*_21_ = *g*_21_, …, *G*_*M* 1_ = *g*_*M* 1_). The prevalence of PSVs by gene can sometimes be obtained from large cohort studies. For example, Anglian Breast Cancer (ABC) Study reported the allele frequencies of BRCA1/BRCA2 genes [26] and Consortium of Investigators of Modifiers of BRCA (CIMBA) reported the frequencies of bins (groups of PSVs) in the BRCA1/BRCA2 genes [21]. Bins were constructed to have equal numbers of PSV carriers with bin length defined by distance in base pairs.

Variant-specific penetrances are often more challenging to estimate because they require sufficiently large sample sizes. Recently, for example, Chen et al. [25] derived equations to estimate the bin-specific penetrance by using the gene-specific PSV penetrance, hazard ratio of breast and ovarian cancer for each bins and frequencies of bins in the BRCA1/BRCA2 genes. Their approach assumed that the gene-specific PSV penetrance is a mixture of bin-specific penetrances. For the application of our variant-specific Mendelian risk prediction model, we grouped the bins and estimated the region-specific penetrances. Section 2.5 illustrates this extension in detail.

### 2.4 Evaluation metrics

To evaluate model performance, we calculated the calibration (O/E), discrimination (AUC), and accuracy (Root Brier score) [27] of the models. For calibration (O/E), we considered the number of observed PSV carriers divided by the number of expected PSV carriers. O/E for a PSV *g*_*m*_ in gene *m* is calculated as

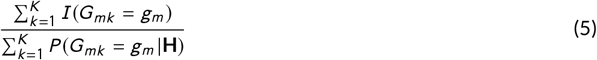

for *k* = 1, …, *K* counselees. Therefore, an O/E of 1 is perfect, calibration higher than 1 means predicted PSV carrier probabilities are on average too low, and lower than 1 means the opposite.

For discrimination we considered, as customary, the AUC. Let *K* ^1^ indicate the set of PSV carriers among counselees and *K* ^0^ indicate the set of non-carriers among counselees. AUC for a PSV *g*_*m*_ in gene *m* is calculated as

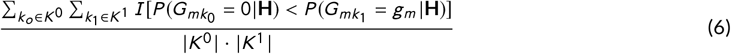

for counselee *k*_1_ ∈ *K* ^1^ who carries PSV *g*_*m*_ and non-carrier counselee *k*_0_ ∈ *K* ^0^. When the AUC of the predicted probabilities for PSV carriers and non-carriers are the same, discrimination is expected to be 0.5.

Accuracy (Root Brier score) was defined as the root mean squared error of the predicted PSV carrier probabilities when compared to the true outcomes. Root Brier score for PSV *g*_*m*_ in gene *m* is calculated as

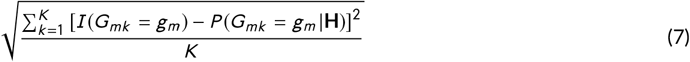

for *k* = 1, …, *K* counselees. Therefore, a Root Brier score of 0 corresponds to a perfect prediction.

### 2.5 Illustration of Proposed Variant-specific Models: BRCAPRO-variant and PanelPRO-variant

BRCA1 gene is a tumor suppressor gene located on chromosome 17q21. BRCA2 gene is a tumor suppressor gene located on chromosome 13q12.3. PSVs in these genes confer increased risk for breast, ovarian, and other cancers [28]. The BRCAPRO model is a Mendelian risk prediction model that incorporates the individual’s family history of breast and ovarian cancer to calculate the gene-specific PSV carrier probability for BRCA1 and BRCA2 genes as well as future risk of breast and ovarian cancer [19]. The BRCAPRO model is widely used in clinics to assess the risk of breast and ovarian cancers; The American Cancer Society (ACS) recommends using BRCAPRO for risk assessment of breast and ovarian cancers [29] along with the Claus [30] and Tyrer-Cuzick models [31]. The PanelPRO [32] model is an extension of the BRCAPRO model, calculating the carrier probability for 22 genes and future risk of 17 cancers.

Due to the sparsity of BRCA1/2 variant-specific risk information, we decided to group PSVs together according to their corresponding cancer risk and calculate the region-specific PSV carrier probability. We propose two model specific extensions; BRCAPRO-variant and PanelPRO-variant models to incorporate variant-specific information through region classifications. The proposed BRCAPRO-variant model incorporates region-specific PSV (bins of PSVs by base pair location) penetrance estimates. The BRCAPRO-variant model is developed to be compatible with BRCAPRO and to be easily incorporated into existing BRCAPRO software. Similarly the proposed PanelPRO-variant model incorporates region-specific PSV penetrance estimates. PanelPRO-variant is compatible with PanelPRO and can be easily incorporated into existing PanelPRO software. For comparison purposes, we ran PanelPRO on solely 2 genes and 2 cancers (namely BRCA1/BRCA2 and Breast/Ovarian).

Rebbeck et al. [21] proposed bins of PSVs by base pair location to identify regions of BRCA1 or BRCA2 genes that are differently associated with breast and ovarian cancer risks. We categorized the bins in the BRCA1 and BRCA2 genes into three regions: the breast cancer clustering region (BCCR) which has significantly elevated risk of breast cancer compared to ovarian cancer, the ovarian cancer clustering region (OCCR) which has significantly elevated risk of ovarian cancer compared to breast cancer, and the no-BCCR & no-OCCR region (“other”), as reported in Rebbeck et al. [21]. We assumed an individual can only belong to a single bin as compound heterozygous PSVs are rare in BRCA genes [33].

The BCCR region in BRCA1 consists of PSVs in nucleotide positions c.179 through c.505, c.4328 through c.4945 and c.5261 through c.5563. The OCCR region in BRCA1 consists of PSVs in nucleotide position c.1380 through c.1674, c.1893 through c.2475, c.2686 through c.3254 and c.4017 through c.4062. The BCCR region in BRCA2 consists of PSVs in nucleotide position c.1 through c.596, c.772 through c.1806 and c.7934 through c.8904. The OCCR region in BRCA2 consists of PSVs in nucleotide position c.3249 through c.5681, c.5946, and c.6645 through c.7471. The “other” region in both BRCA1 and BRCA2 consists of PSVs other than BCCR/OCCR or variants that were not translated to ClinVar’s nomenclature. Penetrance estimates for female breast and ovarian cancers by these regions were obtained from Chen et al. [25] and are shown in Figure 1. Since Rebbeck et al. [33] do not report variant-specific risk for males in BRCA1 and BRCA2, we used the same penetrance for all male carriers, from Tai [34].

**FIGURE 1.**
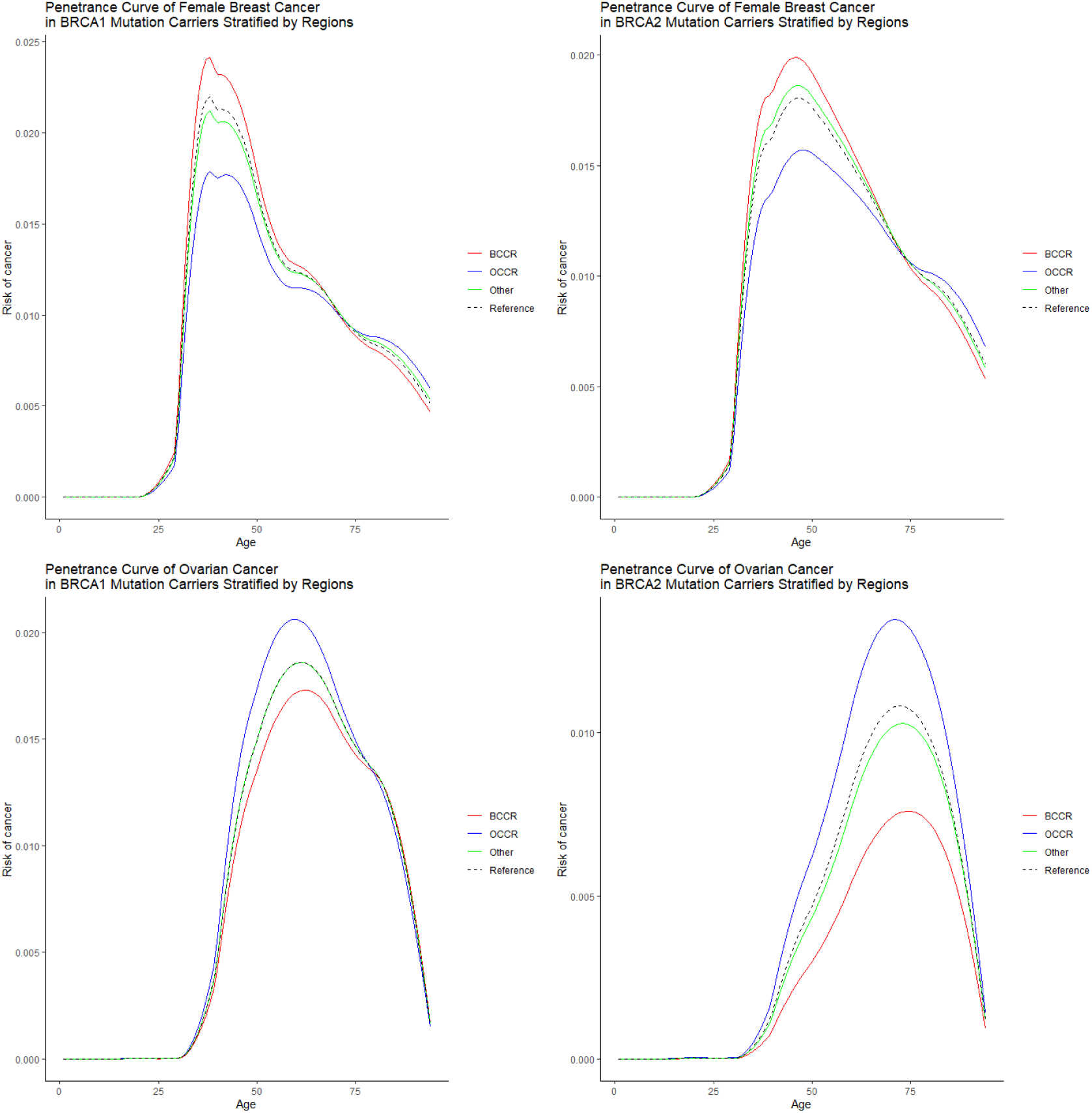
Penetrance functions used in the BRCAPRO-variant and PanelPRO-variant models. Each panel corresponds to a gene-cancer combination. Lines correspond to the BCCR (red), OCCR (blue) and “other” (green) regions as defined in Section 2.5. Dashed curves indicate the general population BRCA1/2 penetrance for females from Chen et al [37].

Our models rely on heavy computations to incorporate all PSVs in any gene. Therefore, we implemented the peeling-paring algorithm [35] in our models to efficiently compute the joint probability of multiple PSVs across genes. Using the peeling-paring algorithm reduces this computation by restricting the maximum number of genes with PSVs. In BRCAPRO-variant and PanelPRO-variant we assumed a maximum two genes with PSVs.

In all models, the allele frequencies of gene-specific PSVs in BRCA1 and BRCA2 were assumed to 0.0005829 and 0.0006760, respectively, which is the general population allele frequencies for Caucasians who are not of Ashkenazi Jewish heritage [26], and 0.006098 and 0.006797, respectively, for Caucasians of Ashkenazi Jewish heritage [36]. In the BRCAPRO-variant and PanelPRO-variant models the proportions of region-specific PSVs in BCCR, OCCR and “other” regions among BRCA1 carriers were assumed to be 0.34, 0.14 and 0.52, respectively [21]. The proportions of region-specific PSVs in BCCR, OCCR and “other” regions among BRCA2 carriers were assumed to be 0.22, 0.34 and 0.44, respectively [21].

Our BRCAPRO-variant and PanelPRO-variant models were developed in the open source R software, version 4.0.2. [38]. Both the models and the code are flexible and can be used for any genes, PSVs, cancers, and model parameters. The BRCAPRO and PanelPRO models are available under the *BayesMendel* package and the *PanelPRO* package, respectively, and can be downloaded from the BayesMendel lab homepage (https://projects.iq.harvard.edu/bayesmendel/our-software). Future package releases will include the BRCAPRO-variant and PanelPRO-variant models.

## 3 SIMULATIONS

### 3.1 Data Generation

Simulations were conducted to evaluate the proposed BRCAPRO-variant model. We generated a general population cohort and a high-risk cohort of families with breast and ovarian cancers. The general population cohort represented a cohort with both underlying and no-underlying hereditary risk, whereas high-risk cohort represented a cohort that meets the NCCN guidelines of genetic/familial high-risk assessment for breast and ovarian cancer [39]. For the general population cohort, PSV status in BCCR, OCCR, and “other” regions in BRCA1/2 for founders of the family were generated based on the allele frequencies of BRCA1/2 PSVs for Caucasians who are not of Ashkenazi Jewish heritage [26] and the frequencies of BCCR, OCCR, and “other” region in BRCA1/BRCA2 genes [21]. The allele frequencies of gene-specific BRCA1/2 PSVs used in this simulation is the one used in the existing BRCAPRO model. PSV carrier statuses were passed to family members based on Mendelian laws of inheritance. Breast and ovarian cancer statuses for 100,000 counselees and their family members were generated based on the PSV carrier statuses and the region-specific penetrances in Figure 1. Breast and ovarian cancer statuses for males were generated based on the penetrance from [34] for all three regions. For our high-risk cohort, PSV carrier status in BCCR, OCCR and “other” regions in BRCA1/2 and the breast and ovarian cancer statuses for 200,000 counselees and their family members were generated based on the same parameters as in the simulated general population cohort, but only counselees who were diagnosed with breast cancer before age 45 were included which reflects the one of the NCCN criteria of Genetic/Familial High-Risk Assessment: Breast and Ovarian [39], resulting in a cohort with 1,966 counselees. Age at death was generated from a normal distribution with mean 85, standard deviation 10 and truncation at 94 (the maximum age allowed in the BRCAPRO and PanelPRO models). We generated families with four generations, all sharing the same family structure presented in Figure 2. The characteristics of the general population simulation cohort are summarized in Table 1 and the high-risk simulation cohort is summarized in Table 2.

**FIGURE 2.**
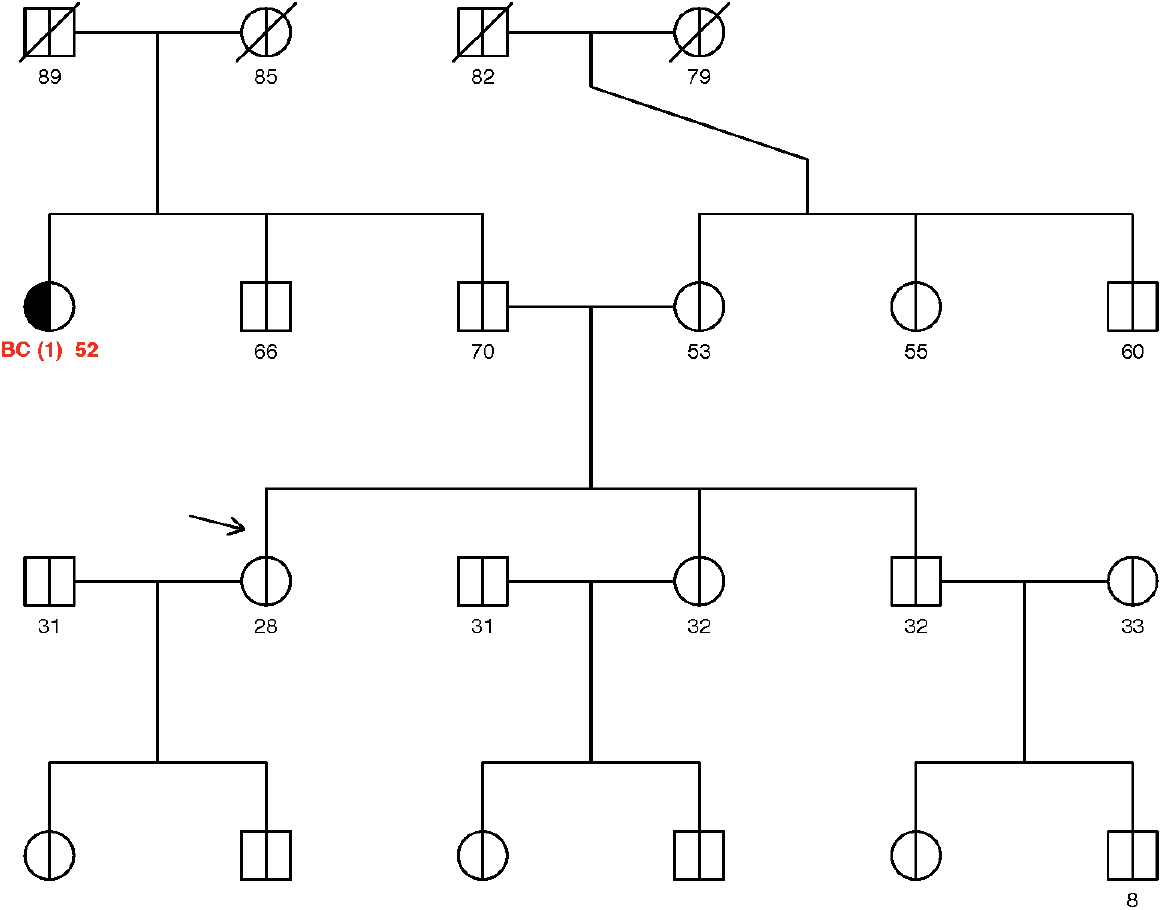
An example simulated pedigree. Family structure includes one paternal aunt and uncle, one maternal aunt and uncle, one sister and one brother for each counselee, and one daughter and one son for each member in the counselee’s generation. The arrow indicates the counselee. Shading on the left indicates that the family member has breast cancer. Shading on the right indicates that the family member has ovarian cancer. A diagonal line indicates that the family member is deceased. Genotypes and phenotypes were generated and varied across families, but family structure was assumed to be the same across all simulated families.

**TABLE 1.**
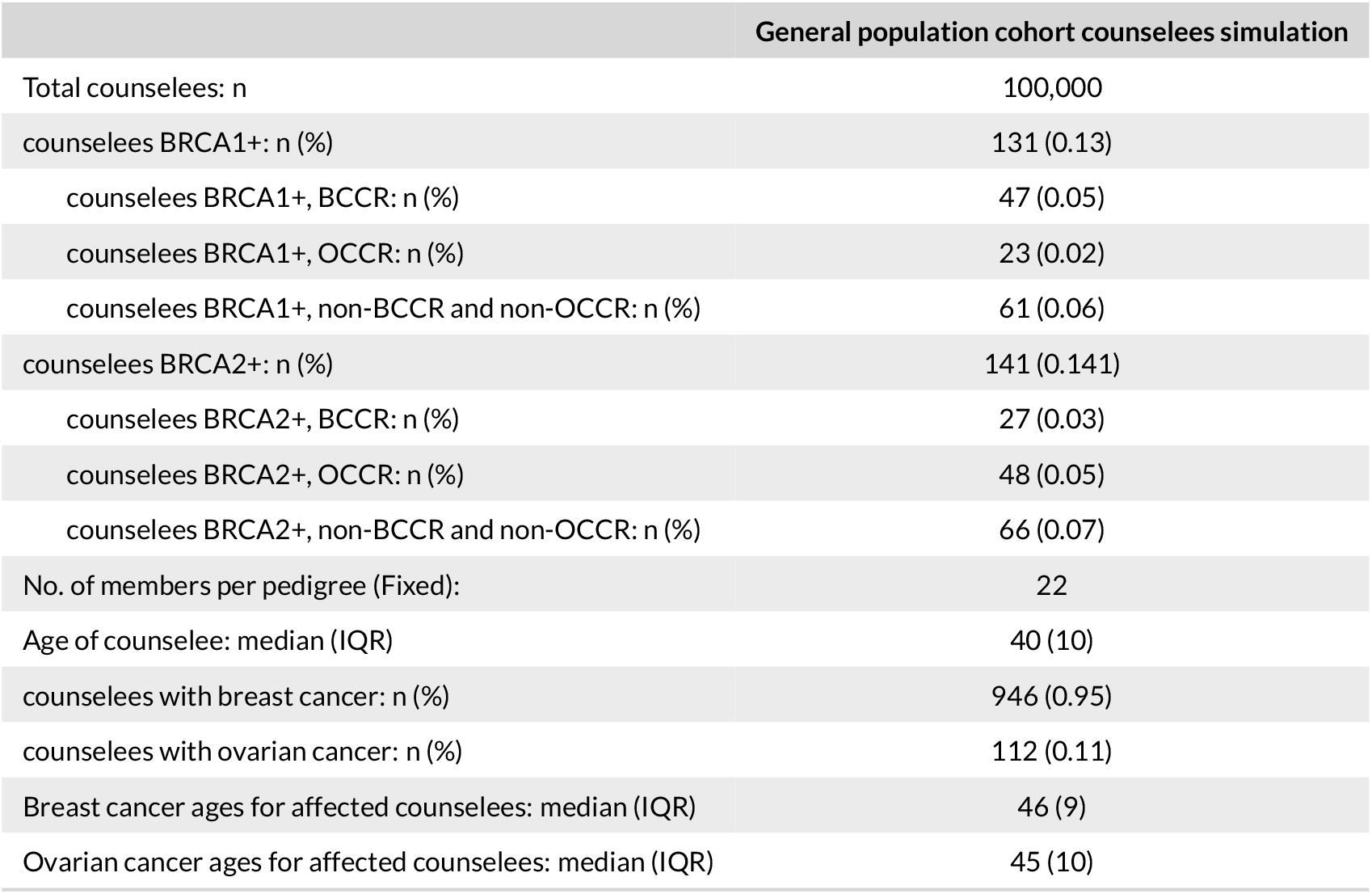
Characteristics of 100,000 simulated general population counselees

**TABLE 2.**
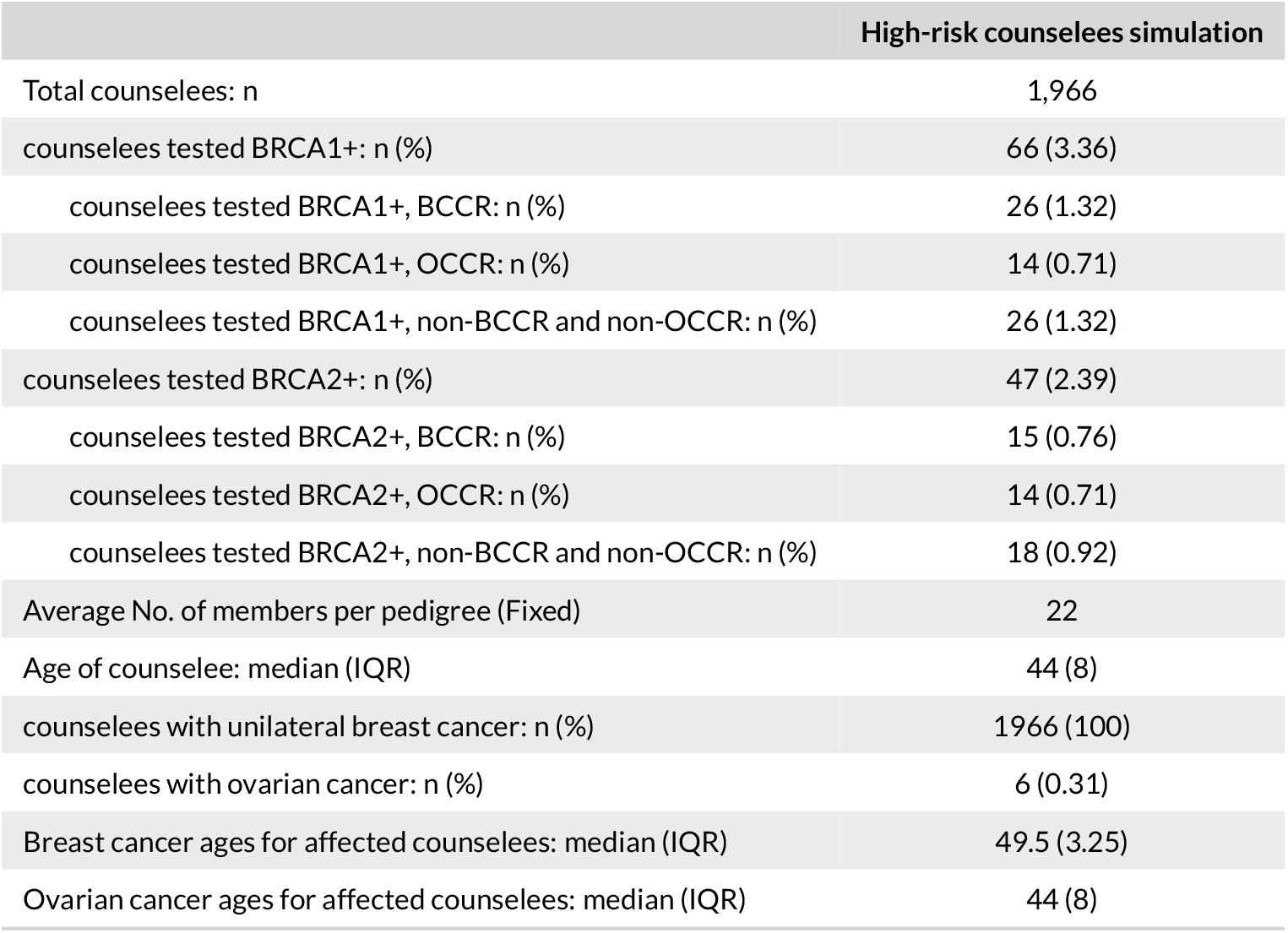
Characteristics of 1,966 simulated high-risk counselees

### 3.2 Simulation results

We evaluated BRCAPRO and the proposed BRCAPRO-variant model based on calibration, discrimination and accuracy of predicting BRCA1 and BRCA2 gene-specific PSV carrier status. Also, we evaluated the BRCAPRO-variant model for prediction of region-specific carrier status in BCCR, OCCR, and “other” regions in the BRCA1 and BRCA2 genes. We calculated confidence intervals using bootstrap replicates. Simulations results are presented in Tables 3 and 4. As the data were generated with variant-specific parameters, we expected that the BRCAPRO-variant model outperforms the BRCAPRO model and we were interested in quantifying the extent of this improvement.

**TABLE 3.** Metrics of model performance, evaluated using 100,000 simulated general population counselees. Confidence intervals were calculated using 100 bootstrap replicates.

|  | O/E | AUC | Root Brier score |
| --- | --- | --- | --- |
| BRCA PSV probability |  |  |  |
| BRCAPRO-variant | 1.06 (0.96, 1.16) | 0.90 (0.89, 0.92) | 0.048 (0.045, 0.050) |
| BRCAPRO | 1.16 (1.05, 1.27) | 0.90 (0.88, 0.92) | 0.048 (0.045, 0.050) |
| BRCA1 PSV probability |  |  |  |
| BRCAPRO-variant | 1.09 (0.93, 1.26) | 0.92 (0.90, 0.95) | 0.034 (0.031, 0.036) |
| BRCAPRO | 1.15 (0.99, 1.34) | 0.92 (0.90, 0.95) | 0.033 (0.031, 0.036) |
| BRCA1, BCCR region-specific PSV probability |  |  |  |
| BRCAPRO-variant | 1.14 (0.81, 1.41) | 0.91 (0.85, 0.96) | 0.021 (0.018, 0.024) |
| BRCA1, OCCR region-specific PSV probability |  |  |  |
| BRCAPRO-variant | 1.37 (0.96, 1.86) | 0.94 (0.89, 0.98) | 0.015 (0.012, 0.018) |
| BRCA1, "other" region-specific PSV probability |  |  |  |
| BRCAPRO-variant | 0.97 (0.75, 1.25) | 0.93 (0.89, 0.97) | 0.023 (0.02, 0.026) |
| BRCA2 PSV probability |  |  |  |
| BRCAPRO-variant | 1.04 (0.87, 1.19) | 0.88 (0.85, 0.91) | 0.037 (0.034, 0.039) |
| BRCAPRO | 1.17 (0.98, 1.34) | 0.88 (0.86, 0.91) | 0.037 (0.034, 0.039) |
| BRCA2, BCCR region-specific PSV probability |  |  |  |
| BRCAPRO-variant | 0.91 (0.63, 1.22) | 0.88 (0.78, 0.94) | 0.016 (0.013, 0.019) |
| BRCA2, OCCR region-specific PSV probability |  |  |  |
| BRCAPRO-variant | 1.03 (0.73, 1.40) | 0.84 (0.78, 0.89) | 0.022 (0.019, 0.025) |
| BRCA2, "other" region-specific PSV probability |  |  |  |
| BRCAPRO-variant | 1.1 (0.87, 1.32) | 0.91 (0.88, 0.94) | 0.025 (0.022, 0.028) |

**TABLE 4.**
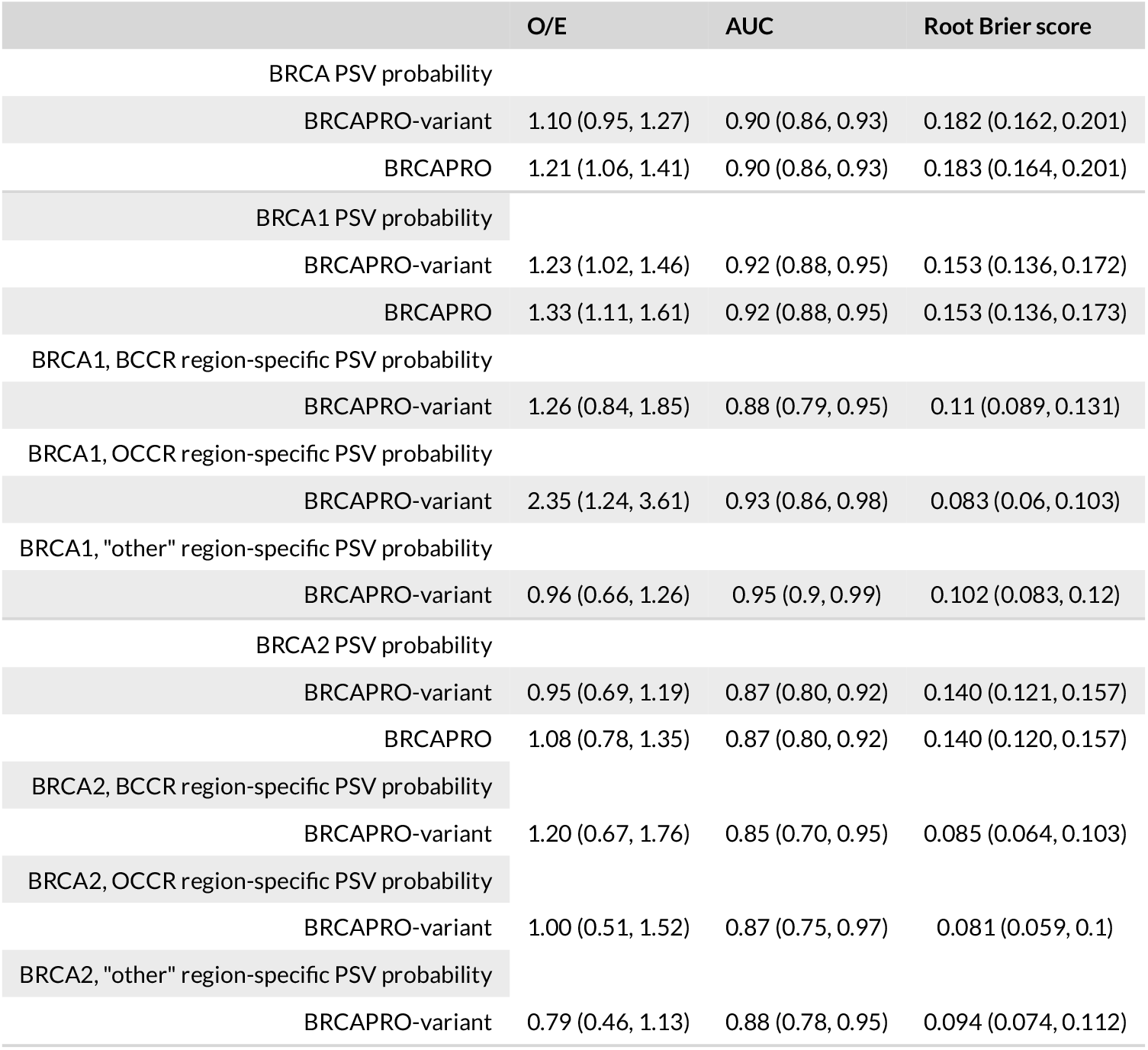
Model performance, evaluated using of 1,966 simulated high-risk counselees. Confidence intervals were calculated using 200 bootstrap replicates.

|  | O/E | AUC | Root Brier score |
| --- | --- | --- | --- |
| BRCA PSV probability |  |  |  |
| BRCAPRO-variant | 1.10 (0.95, 1.27) | 0.90 (0.86, 0.93) | 0.182 (0.162, 0.201) |
| BRCAPRO | 1.21 (1.06, 1.41) | 0.90 (0.86, 0.93) | 0.183 (0.164, 0.201) |
| BRCA1 PSV probability |  |  |  |
| BRCAPRO-variant | 1.23 (1.02, 1.46) | 0.92 (0.88, 0.95) | 0.153 (0.136, 0.172) |
| BRCAPRO | 1.33 (1.11, 1.61) | 0.92 (0.88, 0.95) | 0.153 (0.136, 0.173) |
| BRCA1, BCCR region-specific PSV probability |  |  |  |
| BRCAPRO-variant | 1.26 (0.84, 1.85) | 0.88 (0.79, 0.95) | 0.11 (0.089, 0.131) |
| BRCA1, OCCR region-specific PSV probability |  |  |  |
| BRCAPRO-variant | 2.35 (1.24, 3.61) | 0.93 (0.86, 0.98) | 0.083 (0.06, 0.103) |
| BRCA1, "other" region-specific PSV probability |  |  |  |
| BRCAPRO-variant | 0.96 (0.66, 1.26) | 0.95 (0.9, 0.99) | 0.102 (0.083, 0.12) |
| BRCA2 PSV probability |  |  |  |
| BRCAPRO-variant | 0.95 (0.69, 1.19) | 0.87 (0.80, 0.92) | 0.140 (0.121, 0.157) |
| BRCAPRO | 1.08 (0.78, 1.35) | 0.87 (0.80, 0.92) | 0.140 (0.120, 0.157) |
| BRCA2, BCCR region-specific PSV probability |  |  |  |
| BRCAPRO-variant | 1.20 (0.67, 1.76) | 0.85 (0.70, 0.95) | 0.085 (0.064, 0.103) |
| BRCA2, OCCR region-specific PSV probability |  |  |  |
| BRCAPRO-variant | 1.00 (0.51, 1.52) | 0.87 (0.75, 0.97) | 0.081 (0.059, 0.1) |
| BRCA2, "other" region-specific PSV probability |  |  |  |
| BRCAPRO-variant | 0.79 (0.46, 1.13) | 0.88 (0.78, 0.95) | 0.094 (0.074, 0.112) |

#### 3.2.1 General population cohort simulation results

Table 3 summarizes the general population cohort simulation results. On the region-specific level, the BRCAPRO-variant model was well-calibrated for all region-specific PSVs with O/E = 1.14 (0.81, 1.41) for BRCA1-BCCR, O/E = 1.37 (0.96, 1.86) for BRCA1-OCCR, O/E = 0.97 (0.75, 1.25) for BRCA1-”other”, O/E = 0.91 (0.63, 1.22) for BRCA2-BCCR, O/E = 1.03 (0.73, 1.40) for BRCA2-OCCR, and O/E = 1.1 (0.87, 1.32) for BRCA2-”other”. Imperfect calibration was due to the censoring imposed in the data generation reflecting the life expectancy. The BRCAPRO-variant model discriminated extremely well between region-specific PSVs and non-carriers with AUC = 0.91 (0.85, 0.96) for BRCA1-BCCR, AUC = 0.94 (0.89, 0.98) for BRCA1-OCCR, AUC = 0.93 (0.89, 0.97) for BRCA1-”other”, AUC = 0.88 (0.78, 0.94) for BRCA2-BCCR, AUC = 0.84 (0.78, 0.89) for BRCA2-OCCR, and AUC = 0.91 (0.88, 0.94) for BRCA2-”other”. Also, the BRCAPRO-variant model showed high accuracy for all region-specific PSVs with Root Brier score equal to 0.021 (0.018, 0.024) for BRCA1-BCCR, 0.015 (0.012, 0.018) for BRCA1-OCCR, 0.023 (0.02, 0.026) for BRCA1-”other”, 0.016 (0.013, 0.019) for BRCA2-BCCR, 0.022 (0.019, 0.025) for BRCA2-OCCR, and 0.025 (0.022, 0.028) for BRCA2-”other”. On the gene-specific level, the BRCAPRO-variant model improved the calibration for BRCA (BRCA1+BRCA2) PSVs as well as for BRCA1 PSVs and BRCA2 PSVs separately with the O/E improving from 1.16 (1.05, 1.27) in the BRCAPRO model to 1.06 (0.96, 1.16) in the BRCAPRO-variant model for BRCA PSVs, from 1.15 (0.99, 1.34) to 1.09 (0.93, 1.26) for BRCA1 PSVs, and from 1.17 (0.98, 1.34) to 1.04 (0.87, 1.19) for BRCA2 PSVs. Both the BRCAPRO-variant and the BRCAPRO models showed similar discrimination. Both models showed similar accuracy for BRCA PSVs.

#### 3.2.2 High-risk cohort simulation results

Table 4 summarizes the high-risk cohort simulation results. On the region-specific level, the BRCAPRO-variant model was well-calibrated for BRCA1-BCCR with O/E = 1.26 (0.84, 1.85), BRCA1-”other” with O/E = 0.96 (0.66, 1.26), BRCA2-BCCR with O/E = 1.20 (0.67, 1.76), BRCA2-OCCR with O/E = 1.00 (0.51, 1.52), and BRCA2-”other” with O/E = 0.79 (0.46, 1.13). The BRCAPRO-variant model under-predicted region-specific PSV probabilities for the BRCA1-OCCR region with O/E = 2.35 (1.24, 3.61). This was due to the higher number of BRCA1-OCCR carriers (N=14) than expected number of carriers (N=9.24). The BRCAPRO-variant model discriminated well between region-specific PSV and non-carriers with AUC = 0.88 (0.79, 0.95) for BRCA1-BCCR, AUC = 0.93 (0.86, 0.98) for BRCA1-OCCR, AUC = 0.95 (0.9, 0.99) for BRCA1-”other”, AUC = 0.85 (0.70, 0.95) for BRCA2-BCCR, AUC = 0.87 (0.75, 0.97) for BRCA2-OCCR, and AUC = 0.88 (0.78, 0.95) for BRCA2-”other”. The BRCAPRO-variant model was less accurate in the high-risk population likely because family histories are more similar to each other and therefore it is more difficult to predict the region-specific PSV status, with Root Brier score 0.11 (0.089, 0.131) for BRCA1-BCCR, 0.083 (0.06, 0.103) for BRCA1-OCCR, 0.102 (0.083, 0.12) for BRCA1-”other”, 0.085 (0.064, 0.103) for BRCA2-BCCR, 0.081 (0.059, 0.1) for BRCA2-OCCR, and 0.094 (0.074, 0.112) for BRCA2-”other”. The BRCAPRO-variant model improved the calibration for BRCA (BRCA1+BRCA2) PSVs as well as for BRCA1 PSVs and BRCA2 PSVs, with O/E improving for BRCA PSVs from 1.21 (1.06, 1.41) in the BRCAPRO model to 1.10 (0.95, 1.27) in the BRCAPRO-variant model, for BRCA1 PSVs from 1.33 (1,11, 1.61) to 1.23 (1.02, 1.46), and for BRCA2 PSVs from 1.08 (0.78, 1.35) to 0.95 (0.69, 1.19). Both the BRCAPRO-variant and BRCAPRO models showed similar discrimination and accuracy for BRCA PSVs.

## 4 DATA APPLICATION

### 4.1 Data

We evaluated the BRCAPRO-region, BRCAPRO, PanelPRO-region and PanelPRO models on 1,961 families from the Cancer Genetics Network (CGN) cohort [40]. The CGN cohort is comprised of pedigrees collected in eight high-risk counseling clinics including Huntsman, UT Southwestern, MD Anderson, Baylor, Johns Hopkins, Duke, Penn and Georgetown. The characteristics of the CGN cohort are described in Table 5. The CGN cohort is comprised of high risk counselees such that among the 1,961 counselees, 1,270 (64.8%) counselees had unilateral breast cancer and 162 (8.3%) counselees had ovarian cancer, compared to the 0.95% and 0.11% in the simulated general population cohort and 100% and 0.31% in the simulated high-risk population cohort. 288 (14.7%) counselees were BRCA1 PSV carriers and 136 (6.9%) counselees were BRCA2 PSV carriers. Region-specific PSVs were classified as BCCR, OCCR and “other” based on Section 2.5. There were 175, 37 and 96 BRCA1-BCCR, BRCA1-OCCR and BRCA1-”other” region-specific PSV carriers, respectively, and 24, 33, and 88 BRCA2-BCCR, BRCA2-OCCR and BRCA2-”other” region-specific PSV carriers, respectively.

**TABLE 5.** Characteristics of the Cancer Genetics Network (CGN) cohort

|  | CGN cohorts |
| --- | --- |
| Total counselees: n | 1,961 |
| counselees tested BRCA1+: n (%) | 288 (14.7) |
| counselees tested BRCA1+, BCCR: n (%) | 175 (8.9) |
| counselees tested BRCA1+, OCCR: n (%) | 37 (1.9) |
| counselees tested BRCA1+, non-BCCR and non-OCCR: n (%) | 96 (4.9) |
| counselees tested BRCA2+: n (%) | 136 (6.9) |
| counselees tested BRCA2+, BCCR: n (%) | 24 (1.2) |
| counselees tested BRCA2+, OCCR: n (%) | 33 (1.7) |
| counselees tested BRCA2+, non-BCCR and non-OCCR: n (%) | 88 (4.5) |
| Average No. of members per pedigree (IQR) | 18.5 (9) |
| Age of counselee: median (IQR) | 51.5 (16) |
| counselees with unilateral breast cancer: n (%) | 1270 (64.8) |
| counselees with ovarian cancer: n (%) | 162 (8.3) |
| Breast cancer ages for affected counselees: median (IQR) | 46 (15) |
| Ovarian cancer ages for affected counselees: median (IQR) | 50 (15) |

### 4.2 Results

The BRCAPRO-variant, BRCAPRO, PanelPRO-variant and PanelPRO models were evaluated on the CGN cohort based on calibration, discrimination and accuracy of predicting BRCA1 and BRCA2 gene-specific PSVs probabilities. Also, the BRCAPRO-variant and PanelPRO-variant models were evaluated by predicting region-specific PSVs in the BCCR, OCCR, and “other” regions in the BRCA1 and BRCA2 genes. Confidence intervals were calculated using 200 bootstrap replicates. Bootstrapping in the data application was less computationally intensive compared to the general population simulation data, so 200 bootstrap replicates were used. Results of the models’ performance are presented in Table 6.

**TABLE 6.** Model performance, evaluated on the Cancer Genetics Network (CGN) cohort. Confidence intervals were calculated using 200 bootstrap replicates.

|  | O/E | AUC | Root Brier score |
| --- | --- | --- | --- |
| BRCA PSV probability |  |  |  |
| BRCAPRO-variant | 1.04 (0.97, 1.12) | 0.77 (0.74, 0.79) | 0.390 (0.375, 0.406) |
| BRCAPRO | 1.03 (0.95, 1.10) | 0.77 (0.74, 0.79) | 0.389 (0.372, 0.405) |
| PanelPRO-variant | 0.973 (0.906, 1.048) | 0.767 (0.741, 0.793) | 0.387 (0.372, 0.403) |
| PanelPRO | 0.974 (0.906, 1.049) | 0.767 (0.741, 0.793) | 0.387 (0.372, 0.403) |
| BRCA1 PSV probability |  |  |  |
| BRCAPRO-variant | 1.21 (1.07, 1.33) | 0.80 (0.77, 0.82) | 0.327 (0.311, 0.343) |
| BRCAPRO | 1.16 (1.03, 1.27) | 0.80 (0.77, 0.82) | 0.327 (0.311, 0.343) |
| PanelPRO-variant | 1.127 (1.004, 1.253) | 0.797 (0.767, 0.83) | 0.319 (0.303, 0.338) |
| PanelPRO | 1.169 (1.04, 1.283) | 0.795 (0.766, 0.824) | 0.326 (0.309, 0.343) |
| BRCA1, BCCR region-specific PSV probability |  |  |  |
| BRCAPRO-variant | 1.95 (1.7, 2.25) | 0.78 (0.75, 0.82) | 0.277 (0.258, 0.296) |
| PanelPRO-variant | 1.99 (1.7, 2.32) | 0.79 (0.76, 0.83) | 0.277 (0.258, 0.295) |
| BRCA1, OCCR region-specific PSV probability |  |  |  |
| BRCAPRO-variant | 1.28 (0.88, 1.73) | 0.81 (0.72, 0.87) | 0.137 (0.116, 0.157) |
| PanelPRO-variant | 1.05 (0.69, 1.44) | 0.8 (0.66, 0.87) | 0.127 (0.106, 0.146) |
| BRCA1, "other" region-specific PSV probability |  |  |  |
| BRCAPRO-variant | 0.8 (0.64, 0.94) | 0.79 (0.74, 0.84) | 0.211 (0.191, 0.226) |
| PanelPRO-variant | 0.68 (0.52, 0.81) | 0.79 (0.74, 0.85) | 0.199 (0.182, 0.214) |
| BRCA2 PSV probability |  |  |  |
| BRCAPRO-variant | 0.82 (0.71, 0.96) | 0.73 (0.69, 0.77) | 0.259 (0.243, 0.277) |
| BRCAPRO | 0.85 (0.73, 1.00) | 0.73 (0.68, 0.77) | 0.256 (0.240, 0.275) |
| PanelPRO-variant | 0.75 (0.641, 0.868) | 0.717 (0.671, 0.757) | 0.259 (0.242, 0.277) |
| PanelPRO | 0.743 (0.647, 0.869) | 0.723 (0.676, 0.76) | 0.259 (0.242, 0.275) |
| BRCA2, BCCR region-specific PSV probability |  |  |  |
| BRCAPRO-variant | 0.55 (0.34, 0.81) | 0.76 (0.65, 0.85) | 0.113 (0.092, 0.133) |
| PanelPRO-variant | 0.45 (0.27, 0.66) | 0.7 (0.57, 0.83) | 0.106 (0.087, 0.127) |
| BRCA2, OCCR region-specific PSV probability |  |  |  |
| BRCAPRO-variant | 0.71 (0.51, 0.95) | 0.70 (0.60, 0.80) | 0.129 (0.113, 0.15) |
| PanelPRO-variant | 0.64 (0.46, 0.83) | 0.70 (0.58, 0.80) | 0.13 (0.112, 0.148) |
| BRCA2, "other" region-specific PSV probability |  |  |  |
| BRCAPRO-variant | 1.15 (0.92, 1.38) | 0.68 (0.63, 0.73) | 0.206 (0.186, 0.224) |
| PanelPRO-variant | 1.06 (0.84, 1.28) | 0.68 (0.62, 0.74) | 0.206 (0.185, 0.244) |

On a gene-specific level, variant-specific Mendelian risk prediction models performed similarly to their gene-specific model counterparts. Regarding the BRCA (BRCA1 + BRCA2) PSVs probability, the discrimination and accuracy of the variant-specific models was equal and the calibration was very similar to that of their gene-specific counterpart. Overall differences in the O/E values between two models were less than 5% (Table 6). For BRCA1, the PanelPRO-variant showed a similar performance to PanelPRO with O/E = 1.127 (1.004, 1.253), AUC = 0.797 (0.767, 0.83), Root Brier score = 0.319 (0.303, 0.338) for PanelPRO-variant compared to O/E = 1.169 (1.04, 1.283), AUC = 0.795 (0.766, 0.824), Root Brier score = 0.326 (0.309, 0.343). BRCAPRO had a similar O/E = 1.16 (1.03, 1.27) to that of BRCAPRO-variant’s O/E = 1.21 (1.07, 1.33). The BRCAPRO-variant and BRCAPRO had identical AUC = 0.8 (0.77, 0.82) and Root Brier score = 0.327 (0.311, 0.343). For BRCA2, PanelPRO-variant had O/E = 0.75 (0.641, 0.868) compared to PanelPRO’s O/E = 0.743 (0.647, 0.869). BRCAPRO had O/E = 0.85 (0.73, 1.00) and BRCAPRO-variant had O/E = 0.82 (0.71, 0.96). All four models had similar discrimination with AUC=0.73 (0.68, 0.77) for BRCAPRO and BRCAPRO-variant, and AUC=0.72 (0.62, 0.76) for PanelPRO and PanelPRO-variant. Finally, all four models achieved identical accuracy with a Root Brier score of 0.26 (0.24, 0.28).

For region-specific PSV carrier probabilities, our two variant-specific Mendelian risk prediction models were able to discriminate well, with AUC>0.79 for all BRCA1 regions and AUC>0.68 for all BRCA2 regions. Accuracy in region-specific PSV carrier probabilities was also high for our two models, with Root Brier scores less than 0.28 for BRCA1 and less than 0.21 for BRCA2. The OCCR region was well calibrated in BRCA1, with O/E=1.05 (0.69, 1.44) in PanelPRO-variant and O/E=1.28 (0.88, 1.73) in BRCAPRO-variant, while the “other” region was well calibrated in BRCA2 with O/E=1.06 (0.84, 1.28) in PanelPRO-variant and O/E=1.15 (0.92, 1.38) in BRCAPRO-variant. We observed unstable calibration of the BCCR region in both BRCA1 and BRCA2. BCCR carrier probabilities were under-predicted in BRCA1, with O/E=1.95 (1.7, 2.25) for BRCAPRO-variant and O/E=1.99 (1.7, 2.32) for PanelPRO-variant. On the other hand, BCCR carrier probabilities were over-predicted in BRCA2 with O/E=0.55 (0.34, 0.81) for BRCAPRO-variant and O/E=0.45 (0.27, 0.66) for PanelPRO-variant. This may be due to the ascertainment criteria for the CGN cohort which is different from the ascertainment criteria for CIMBA cohort which was used to obtain the variant-specific parameters [21]. The frequencies of BCCR, OCCR, and “other” in BRCA1 in CIMBA cohort that was used to estimate the prior probability was 0.34, 0.14, 0.52, whereas the frequencies of BCCR, OCCR, and “other” in BRCA1 in CGN cohort were 0.57, 0.12, and 0.31. Similarly, the frequencies of BCCR, OCCR, and “other” in BRCA2 in CIMBA cohort were 0.22, 0.34, and 0.44, whereas the frequencies of BCCR, OCCR, and “other” in BRCA2 in CGN cohort were 0.17, 0.23, and 0.60. This discrepant frequencies impacted the prior distributions of genotypes and therefore impacted the expected number of region-specific PSV carriers as well as the calibration result. To understand the under-predicted carrier probabilities for the BRCA1 regions and the over-predicted carrier probabilities for the BRCA2 regions, sensitivity analyses were conducted and discussed in Section 5.

## 5 DISCUSSION

Our proposed BRCAPRO-variant and PanelPRO-variant models are the first Mendelian risk prediction models that can provide variant-specific PSV probabilities. The proposed models provide refined cancer risk estimates that may be of value in tailoring cancer genetic counseling, decision making, and prevention strategies. In addition, the BRCAPRO-variant and PanelPRO-variant models provide accurate estimates of region-specific PSV carrier probabilities and future risk of cancers. The lifetime risk of ovarian cancer for a 25-year-old woman who has a PSV in BRCA2 is 39% assuming no censoring, but depending on her PSV, this risk ranges from 27% to 49% (Figure 1) [25].

Simulation study results showed that the BRCAPRO-variant model is well calibrated with satisfactory discrimination and accuracy. We evaluated the BRCAPRO-variant model on a simulated general population / high-risk counselees, albeit the high-risk counselees did not resemble the CGN cohort as the inclusion criteria differs. On the gene-specific level, the BRCAPRO-variant model showed better calibration than the BRCAPRO model with similar discrimination and accuracy. On the region-specific level, the BRCAPRO-variant model showed stable calibration with AUC > 0.84 and Root Brier score < 0.025. In the CGN cohort, PanelPRO-variant and PanelPRO showed similar performance in terms of calibration, discrimination and accuracy, and BRCAPRO-variant and BRCAPRO showed similar performance in calibration, discrimination and accuracy regarding gene-specific PSVs probabilities. The variant-specific Mendelian risk prediction models showed an ability to discriminate between specific region-specific PSVs and non-carriers, with AUC > 0.78 for both BRCAPRO-variant and PanelPRO-variant in BRCA1 OCCR, BCCR and “other” region-specific PSVs and AUC > 0.68 for BRCA2 BCCR, OCCR and “other” region-specific PSVs. This demonstrates the predictive ability that variant-specific Mendelian risk prediction models can achieve for region-specific PSV carrier probabilities, while providing a similar, or sometimes improved, discriminative ability to traditional Mendelian risk prediction models for gene-specific carrier probabilities. Our BRCAPRO-variant and PanelPRO-variant models can provide clinically relevant risk scores that can help distinguish breast and ovarian cancer risks and thus users can refine patients’ cancer prevention strategies. We also expect that the model will have a better ability to recognize inherited susceptibility that manifests itself mostly through breast or mostly through ovarian cancer phenotypes.

**TABLE S1.** Number of region-specific PSV carriers in the Cancer Genetics Network (CGN) cohort among each scenario considered for the sensitivity analysis

|  | BCCR | OCCR | Other |
| --- | --- | --- | --- |
| BRCA1+ |  |  |  |
| P-value | 175 | 37 | 96 |
| Effect-size | 199 | 81 | 28 |
| Contiguity | 192 | 88 | 28 |
| BRCA2+ |  |  |  |
| P-value | 24 | 33 | 88 |
| Effect-size | 99 | 40 | 6 |
| Contiguity | 51 | 88 | 6 |

We recognize that variant-specific Mendelian risk prediction models have the potential to continuously improve as new data becomes available. Indeed the BRCAPRO-region and PanelPRO-region models showed unstable calibration on region-specific level in the CGN cohort indicating an under-prediction of the BRCA1-BCCR carriers (Table 6) and an over-prediction of the BRCA2-BCCR PSV. This may be attributed to the number of PSVs in each region greatly varying based on how regions are defined and how the PSVs in the CGN cohort are classified (based on Rebbeck et al) [21]. We conducted additional sensitivity analyses to investigate this unstable calibration, where we see that depending on our region classification (effect-size, p-value and continuous region), the number of carriers varied as shown in Table S1 but region-specific penetrance estimates have very little variability as shown in Figure S1. This led to unstable calibration over the varying scenarios as shown in Table S2. *P* -value classification showed the best accuracy for all region-specific level prediction. Future work is needed to define regions which are clinically meaningful and achieve stable calibration. Our model assumed that prevalence of genes and PSV-specific penetrances are correctly specified. However, this assumption may not always be accurate or testable. The current modeling framework could be extended to incorporate this uncertainty via a further probability distribution, as described in [41].

Variant-specific Mendelian risk prediction models may also be applicable to patients with variants of uncertain significance (VUS). van Marcke et al. report that as many as 23% of patients have VUS [42]. According to the ACMG guidelines, individuals with a VUS should not be managed in the same way as carriers of PSVs [43]. For these individuals, although they have already undergone testing for germline PSVs, pre-test probabilities can still provide useful information, as they indirectly inform on the potential pathogenicity of the VUS. For example, if a counselee has a VUS in the BCCR region of the BRCA1 gene and has high carrier score for carrying a PSV in BCCR, she may benefit from considering additional preventative measures including mammography screening, MRI, hormone replacement treatment, and mastectomy.

**TABLE S2.** Model performance, evaluated on the Cancer Genetics Network (CGN) cohort in all scenarios considered for the sensitivity analyses

|  | O/E | AUC | Root Brier score |
| --- | --- | --- | --- |
| BRCA PSV probability |  |  |  |
| BRCAPRO | 1.03 (0.95, 1.10) | 0.77 (0.74, 0.79) | 0.389 (0.372, 0.405) |
| BRCAPRO-variant (P-value) | 1.04 (0.97, 1.12) | 0.77 (0.74, 0.79) | 0.39 (0.375, 0.406) |
| BRCAPRO-variant (Effect-size) | 1.04 (0.97, 1.12) | 0.77 (0.74, 0.79) | 0.39 (0.375, 0.406) |
| BRCAPRO-variant (Contiguity) | 1.04 (0.97, 1.13) | 0.77 (0.74, 0.79) | 0.39 (0.374, 0.406) |
| BRCA1 PSV probability |  |  |  |
| BRCAPRO | 1.16 (1.03, 1.27) | 0.80 (0.77, 0.82) | 0.327 (0.311, 0.343) |
| BRCAPRO-variant (P-value) | 1.21 (1.07, 1.33) | 0.8 (0.77, 0.82) | 0.327 (0.311, 0.343) |
| BRCAPRO-variant (Effect-size) | 1.21 (1.07, 1.32) | 0.8 (0.77, 0.82) | 0.327 (0.311, 0.343) |
| BRCAPRO-variant (Contiguity) | 1.21 (1.07, 1.32) | 0.8 (0.77, 0.82) | 0.327 (0.311, 0.344) |
| BRCA1, BCCR region-specific PSV probability |  |  |  |
| BRCAPRO-variant (P-value) | 1.95 (1.70, 2.25) | 0.78 (0.75, 0.82) | 0.277 (0.258, 0.296) |
| BRCAPRO-variant (Effect-size) | 1.56 (1.36, 1.77) | 0.8 (0.76, 0.83) | 0.29 (0.272, 0.309) |
| BRCAPRO-variant (Contiguity) | 1.26 (1.1, 1.42) | 0.8 (0.76, 0.83) | 0.286 (0.269, 0.303) |
| BRCA1, OCCR region-specific PSV probability |  |  |  |
| BRCAPRO-variant (P-value) | 1.28 (0.88, 1.73) | 0.81 (0.72, 0.87) | 0.137 (0.116, 0.157) |
| BRCAPRO-variant (Effect-size) | 0.76 (0.62, 0.92) | 0.81 (0.77, 0.87) | 0.203 (0.187, 0.218) |
| BRCAPRO-variant (Contiguity) | 1.06 (0.86, 1.27) | 0.81 (0.76, 0.86) | 0.203 (0.185, 0.22) |
| BRCA1, "other" region-specific PSV probability |  |  |  |
| BRCAPRO-variant (P-value) | 0.8 (0.64, 0.94) | 0.79 (0.74, 0.84) | 0.211 (0.191, 0.226) |
| BRCAPRO-variant (Effect-size) |  | Not Defined |  |
| BRCAPRO-variant (Contiguity) |  | Not Defined |  |
| BRCA2 PSV probability |  |  |  |
| BRCAPRO | 0.85 (0.73, 1.00) | 0.73 (0.68, 0.77) | 0.256 (0.240, 0.275) |
| BRCAPRO-variant (P-value) | 0.82 (0.71, 0.96) | 0.73 (0.69, 0.77) | 0.259 (0.243, 0.277) |
| BRCAPRO-variant (Effect-size) | 0.82 (0.71, 0.96) | 0.73 (0.69, 0.77) | 0.259 (0.243, 0.277) |
| BRCAPRO-variant (Contiguity) | 0.83 (0.71, 0.98) | 0.73 (0.69, 0.77) | 0.258 (0.242, 0.276) |
| BRCA2, BCCR region-specific PSV probability |  |  |  |
| BRCAPRO-variant (P-value) | 0.55 (0.34, 0.81) | 0.76 (0.65, 0.85) | 0.113 (0.092, 0.133) |
| BRCAPRO-variant (Effect-size) | 0.95 (0.77, 1.16) | 0.71 (0.66, 0.76) | 0.218 (0.202, 0.237) |
| BRCAPRO-variant (Contiguity) | 0.54 (0.4, 0.71) | 0.74 (0.65, 0.81) | 0.166 (0.149, 0.184) |
| BRCA2, OCCR region-specific PSV probability |  |  |  |
| BRCAPRO-variant (P-value) | 0.71 (0.51, 0.95) | 0.70 (0.60, 0.80) | 0.129 (0.113, 0.150) |
| BRCAPRO-variant (Effect-size) | 0.65 (0.48, 0.84) | 0.7 (0.61, 0.78) | 0.145 (0.128, 0.162) |
| BRCAPRO-variant (Contiguity) | 1.23 (1.02, 1.46) | 0.69 (0.64, 0.75) | 0.206 (0.186, 0.223) |
| BRCA2, "other" region-specific PSV probability |  |  |  |
| BRCAPRO-variant (P-value) | 1.15 (0.92, 1.38) | 0.68 (0.63, 0.73) | 0.206 (0.186, 0.224) |
| BRCAPRO-variant (Effect-size) |  | Not Defined |  |
| BRCAPRO-variant (Contiguity) |  | Not Defined |  |

We introduced a variant-specific Mendelian risk prediction model that provides variant-specific carrier probabilities to help users optimize risk assessment and cancer prevention strategies. This novel model extension are generalizable to any gene and may have clinical value as our understanding of the genes and PSVs associated with cancer risk become more complete, and our ability to optimize the clinical utility of genetic susceptibility relies increasingly on complex genetic architecture of these genes.

## Acknowledgements

This work is supported by the National Cancer Institute at the National Institutes of Health [R01-CA207365-01A1 and 5P30CA006516-54]. Computations in this paper were run on the Odyssey cluster supported by the FAS Division of Science, Research Computing Group at Harvard University. Data were collected within the framework of the NCI’s Cancer Genetics Network, combining data previously collected at each Cancer Genetics Network center and at City of Hope National Medical Center. Most data predated Cancer Genetics Network activities, but the Cancer Genetics Network provided the venue for the pooled analysis. The authors thank Connie Griffin for her Cancer Genetics Network leadership at the Johns Hopkins University, Kelly Qu for support with database management at Johns Hopkins University, Jihong Zong for collecting and transmitting data at M.D. Anderson Cancer Center, and Neil Malloy for assistance with data coordination at Massachusetts General Hospital.

## Data Availability Statement

De-identified data that support the findings of this study are available on request. The data are not publicly available due to privacy restrictions.

## Abbreviations

DNA: Deoxyribonucleic acid
PSV: Pathogenic sequence variant
BRCA: breast cancer gene
CGN: Cancer Genetics Network
VUS: variants of uncertain significance

## Supplementary Materials

### I. Classification of BCCR/OCCR and “other”

We considered additional approaches to classify the BRCA1/2 PSV into regions as part of a sensitivity analysis. We considered an effect-size criteria, which classifies the BCCR/OCCR based on the hazard ratio.

If the higher hazard ratio of breast cancer was observed than ovarian cancer in CIMBA, then that region was classified as BCCR and OCCR vice versa. Based on effect-size criteria, the BCCR in BRCA1 consisted of PSV in nucleotide position c.1 through c.151, c.178 through c.927, c.3483 through c.3661 and c.4063 through c.5563. The OCCR in BRCA1 consisted of PSV in nucleotide position c.68 through c.69, c.928 through c.3482 and c.3662 through c.4062. The BCCR in BRCA2 consisted of PSV in nucleotide position c.1 through c.596, c.772 through c.3248, c.5682 through c.5945, c.5947 through c.6275 and c.7472 through c.9925. The OCCR in BRCA2 consisted of PSV in nucleotide position c.597 through c.771, c.3249 through c.5681, c.5946 and c.6276 through c.7471. Variant that was not translated to Clinvar’s nomenclature was classified as “other”. We also considered a contiguity criteria which classifies the BCCR/OCCR assuming that BCCR/OCCR is contiguously located.

If the higher hazard ratio of breast cancer was observed than ovarian cancer for consecutive bins in CIMBA, then that region was classified as BCCR and OCCR vice versa. Based on contiguity criteria, the BCCR in BRCA1 consisted of PSV in nucleotide position c.1 through c.927 and c.4063 through c.5563. The OCCR in BRCA1 consisted of PSV in nucleotide position c.928 through c.4062. The BCCR in BRCA2 consisted of PSV in nucleotide position c.1 through c.3248 and c.7472 through c.9925. The OCCR in BRCA2 consisted of PSV in nucleotide position c.3249 through c.7471. Variant that was not translated to Clinvar’s nomenclature was classified as “other”.

**FIGURE S1.**
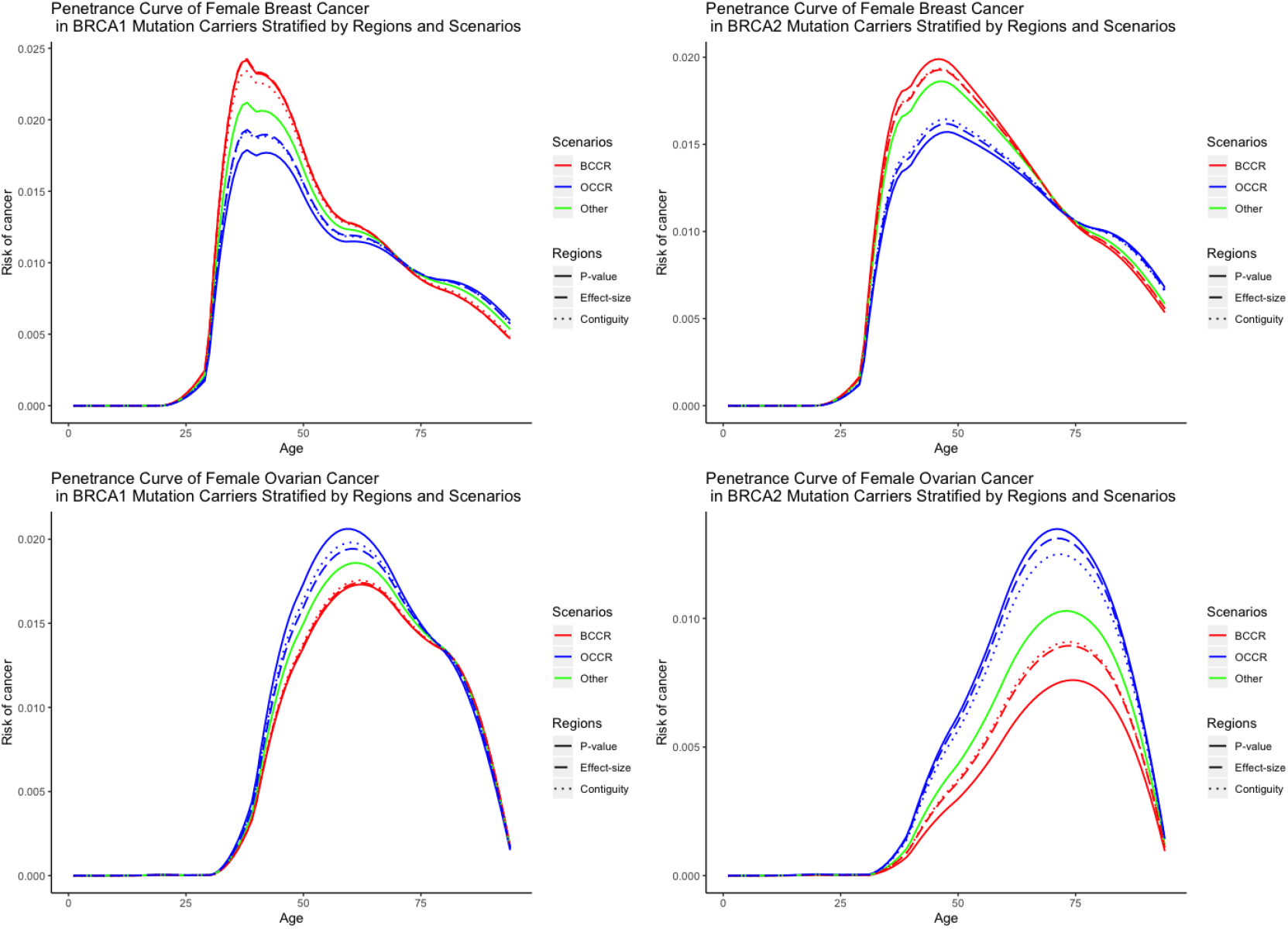
Penetrance curves for sensitivity analysis where various classification scenarios are considered. *P* -value (solid curve) indicates the classification of BCCR/OCCR and “other” based on p-values. Effect-size (long-dash curve) indicates the classification of BCCR and OCCR based on effect sizes. Contiguity (dotted curve) indicates the classification of BCCR and OCCR based on contiguous regions. BCCR (red curve) indicates breast cancer clustering region which has elevated risk of breast cancer compared to ovarian cancer. OCCR (blue curve) indicates ovarian cancer clustering region which has elevated risk of ovarian cancer compared to bresat cancer. “other” (green curve) indicates regions that are not belong to BCCR and OCCR. Note that only P-value classification has the “other” category.

